# Reducing false positives in tractography with microstructural and anatomical priors

**DOI:** 10.1101/608349

**Authors:** Simona Schiavi, Muhamed Barakovic, Mario Ocampo-Pineda, Maxime Descoteaux, Jean-Philippe Thiran, Alessandro Daducci

## Abstract

Tractography is a family of algorithms that use diffusion-weighted magnetic resonance imaging data to reconstruct the white matter pathways of the brain. Although it has been proven to be particularly effective for studying non-invasively the neuronal architecture of the brain, recent studies have highlighted that the large incidence of false positive connections retrieved by these techniques can significantly bias any connectivity analysis. Some solutions have been proposed to overcome this issue and the ones relying on convex optimization framework showed a significant improvement. Here we propose an evolution of the Convex Optimization Modeling for Microstructure Informed Tractography (COMMIT) framework, that combines basic prior knowledge about brain anatomy with group-sparsity regularization into the optimization problem. We show that the new formulation dramatically reduces the incidence of false positives in synthetic DW-MRI data.

## 1 Introduction

Diffusion-weighted magnetic resonance imaging (DW-MRI) is a non-invasive imaging technique that is sensitive to how water molecules diffuse in biological tissues [1]. By capturing coherent orientations of maximal diffusion among neighboring voxels inside white matter (WM), tractography algorithms can infer the trajectories of the major neuronal pathways in the brain [2] and can be used to map the human connectome [3, 4]. A *connectome* is a comprehensive map of the brain’s neuronal connections and is typically represented as a graph, where nodes correspond to gray-matter nuclei and edges to the WM connections between them. Using this representation, brain connectivity can be studied by means of graph theory and network science [5, 6] and, over the years, this approach has been successfully exploited to study a wide range of neurological conditions [7, 8].

Because of the large difference between the resolution of DW-MRI acquisitions (order of millimeters) and the size of axons (order of micrometers), a single tract reconstructed with tractography, called streamline, cannot represent a single axon but rather a coherent set of real anatomical fibers. This discrepancy introduces ambiguities that are difficult to resolve and, indeed, recent studies have highlighted that the accuracy of tractography is inherently limited, raising serious concerns about its use in connectivity analysis. Thomas et al. [9] highlighted that existing methods suffer from an intrinsic trade-off between *sensitivity*, i.e. capability of reconstructing real WM bundles, and *specificity*, i.e. retrieving only true connections, even when using very high quality data. Using graph theory, Zalesky et al. [10] demonstrated that specificity is crucial when using tractography to study the topological properties of brain networks, and twice as important than sensitivity. However, Maier-Hein et al. [11] actually showed that specificity represents the main bottleneck for tractography and tractograms are typically polluted by many false positives. Hence, improving the specificity of tractography, i.e. reducing false positives, represents a major challenge in computational neuroscience towards a more veridical characterization of brain connectivity.

A number of solutions have been proposed recently to *improve the accuracy of tractography reconstructions*, such as MicroTrack [12], Spherical-deconvolution Informed Filtering of Tractograms (SIFT) [13, 14], Linear Fascicle Evaluation (LiFE) [15] and Convex Optimization Modeling for Microstructure Informed Tractography (COMMIT) [16, 17]. The common approach behind these methods consists of combining the reconstructed set of streamlines, i.e. tractogram, with signal forward-models to assess their actual contribution to the acquired DW-MR data and filter out the most implausible using optimization [18]. Despite the tractograms filtered with such techniques provide more biologically accurate estimates of the connectivity [19], none of them has been proven effective in reducing false positives. All methods are purely data-driven and rely only on the acquired DW-MR signal to evaluate the effective contribution of each streamline. In all cases, the estimation process considers all streamlines as independent entities, thus ignoring the fact that in the central nervous system axons are naturally organized in bundles. It is worth noting that this fact is actually at the core of connectome mapping, where streamlines are grouped together in bundles and considered as single edges of the resulting brain network [3, 4].

In this study, we present a novel processing framework for dramatically reducing the incidence of these false positives, i.e. improving specificity, without affecting the true ones, i.e. sensitivity. Tractography algorithms usually exploit only the directional information estimated with DW-MRI: we speculate that this information is not enough and we advocate the need for additional data to help tractography producing more accurate reconstructions. Our formulation injects basic prior knowledge about brain anatomy by adding to the COMMIT framework an efficient regularization term which can help *resolving ambiguous configurations and reducing the false positives*. We evaluated quantitatively the effectiveness of our formulation on different tractography reconstructions using a realistic digital phantom with known ground-truth.

## 2 Materials and methods

### 2.1 Microstructure informed tractography

Given a DW-MR image **I** and a tractogram 𝒯, the acquired data can be seen as **I** = 𝓐(𝒯) + *η*, where 𝓐: *𝒯* → **I** is an operator describing the signal contribution of each fiber to the *n*_*d*_ q-space samples acquired in the *n*_*v*_ = *n*_*x*_*n*_*y*_*n*_*z*_ voxels of 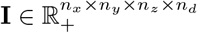 and *η* is the acquisition noise. The goal of tractography is to solve the inverse problem, i.e. finding the set of streamlines 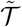 that best describe the acquired image **I**. The term “microstructure informed tractography” refers to a relatively novel area of research [18] whose aim is to obtain more quantitative and biologically meaningful estimates of brain connectivity by complementing tractography with biophysical models of the tissue microstructure [20]. Several solutions have been proposed [12–17] but the originality of the Convex Optimization Modeling for Microstructure Informed Tractography (COMMIT) [16, 17] lies in the possibility to express tractography and tissue microstructure in a unified framework and solve this inverse problem using convex optimization. The signal in each voxel of **I** is described as a linear combination of the diffusion arising from all the fibers of 𝒯 that intersect the voxel, in addition to local contributions from other tissues, e.g. cerebrospinal fluid (CSF). The joint problem can then be expressed as a system of linear equations:

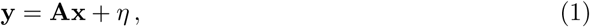

where the vector 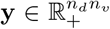 contains the *n*_*d*_ DW-MR measurements acquired in the *n*_*v*_ voxels of **I**, the matrix **A** ∈ ℝ^*n*_*d*_*n*_*v*_ × *n*_*c*_^ encodes the potential contributions of all streamlines in (and possibly other tissues) to the signal in each voxel according to a given multi-compartment model and *η* accounts for both acquisition noise and modeling errors. The positive weights 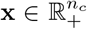 represent the actual contributions of the *n*_*c*_ compartments, encoded as columns of **A**, needed to explain the acquired data **I** and can be estimated using non-negative least squares (NNLS):

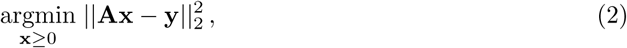

where || · || _2_ is the Euclidean norm in ℝ^*n*^.

Any multi-compartment model [20] can be virtually used in COMMIT. In general, a multi-compartment model assumes different diffusion behaviors according to the microstructure geometrical properties. For neuronal tissue, a common assumption is to distinguish between three compartments [12]: intra-axonal (IA, mimicking the restricted movement of water molecules inside axons), extra-axonal (EA, mimicking the hindered movement outside axons) and isotropic (ISO, mimicking the free movement of the water like in CSF). The linear operator **A** is typically a block matrix of this form:

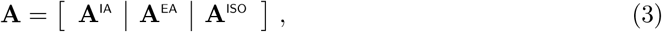

where *n*_*c*_ = *n*_*r*_+*n*_*h*_+*n*_*i*_ and the sub-matrices **A**^IA^ ∈ ℝ^*n*_*d*_*n*_*v*_ × *n*_*r*_^, **A**^EA^ ∈ ℝ^*n*_*d*_*n*_*v*_ × *n*_*h*_^ and **A**^ISO^ ∈ ℝ^*n*_*d*_*n*_*v*_ × *n*_*i*_^ encode, respectively, the *n*_*r*_ restricted, *n*_*h*_ hindered and *n*_*i*_ isotropic contributions to the image.

#### Illustrative toy example

To illustrate this estimation process, let’s consider the synthetic toy example shown in Fig. 1a. In the left panel we display the orientation distribution functions (ODF) simulated in each voxel, which were used to reconstruct the three streamlines visualized in the middle panel using a generic tractography algorithm. The right panel shows the forward model we adopted to construct the operator 𝓐: a *stick* to account for the anisotropic contributions of the streamlines and a *ball* to consider possible CSF contaminations [20]. Fig. 1b illustrate the components of the linear system **y** = **Ax** that we want to solve using COMMIT. In the column vector **y** we concatenate the data simulated in each voxel. The matrix **A** is constructed by first checking which voxels are intersected by the reconstructed streamlines: fiber 1 crosses voxels 1 and 2, fiber 2 crosses voxels 1 and 3; fiber 3 crosses voxel 2, 3 and 4. We then create one column for each streamline and store in the rows corresponding to each voxel it traverses the contribution of a *stick* oriented in the same direction of the streamline; if a streamline does not cross a voxel, the corresponding rows are set to 0. To account for the possible presence of CSF in a voxel we add four columns and, in each of them, we put 0 everywhere except in the rows corresponding to a distinct voxel where we insert an isotropic contribution according to the *ball* model.

**Figure 1:**
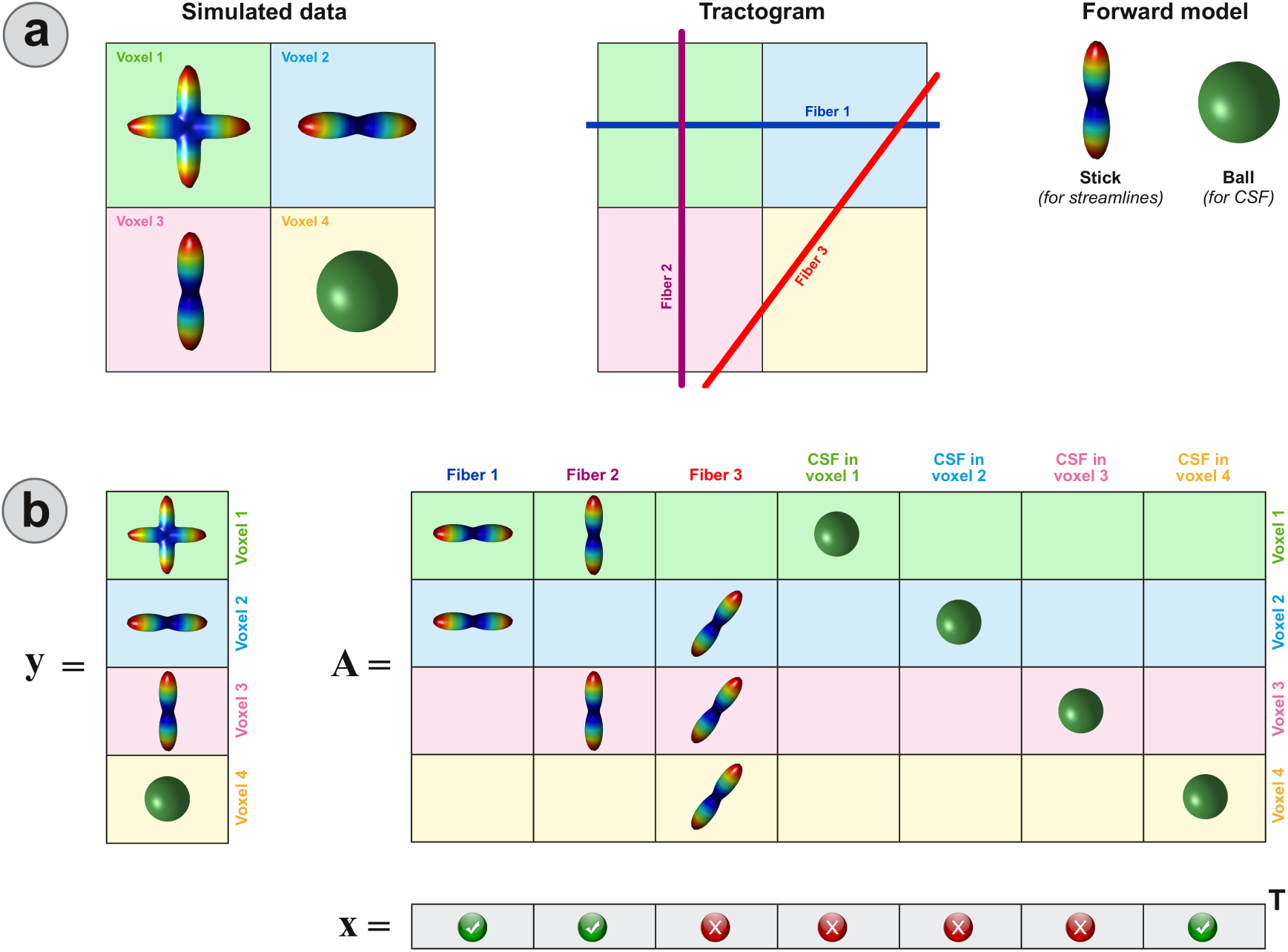
Synthetic toy example to illustrate the modeling and the parameter estimation using the COMMIT framework. (**a**) The simulated orientation distribution functions (ODF), a possible tractogram estimated with a generic tractography algorithm and the forward-model used to associate a signal contribution to each streamline. (**b**) The corresponding vector **y** containing the simulated data in all voxels, the matrix **A** encoding the signal contributions of each streamline according to the chosen forward-model (as well as potential presence of CSF) and the coefficients **x** estimated by COMMIT.

Every column in **A** is controlled by a different contribution in **x** and, for a given configuration of contributions **x**, the predicted signal is obtained by performing the multiplication **Ax**. COMMIT seeks for the optimal configuration of **x**, which must be positive, such that the predicted signal, i.e. **Ax**, is as close as possible to the measured signal, i.e. **y**, hence it tries to minimize their difference, i.e. argmin 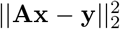. According to matrix-vector multiplication properties, we can immediately notice that to obtain the correct profile in voxel 1, we must have a positive contribution in the first two entries of **x** but 0 in **x**_4_ since there is no CSF in voxel 1. To assign the values to the remaining entries of **x** we continue the multiplication. Looking at voxel 2 we observe that **x**_3_ = **x**_5_ = 0, while from the third and forth voxels we obtain **x**_6_ = 0 and **x**_7_ = 1 respectively. The entries of **x** are uniquely determined and, since **x**_3_ = 0, fiber 3 will be marked as false positive and removed from the tractogram.

### 2.2 Injecting priors about brain anatomy and its organization

The purpose of this study was to evaluate whether we could improve the sensitivity/specificity trade-off of tractography by taking into consideration two very basic observations about WM anatomy during the estimation process: (i) streamlines are not “just lines” but represent neuronal fibers, and (ii) such neuronal fibers are naturally organized in bundles. To enforce the first prior knowledge, we implemented in **A** a simple forward-model that assigns a contribution, i.e. volume or cross-sectional area, to each streamline of the input tractogram 𝒯 proportionally to its length inside each voxel. Then with Eq. 2 we require that the total amount of streamlines that traverse a voxel must sum up to the actual intra-axonal signal fraction in that voxel, which can be estimated in every voxel of the brain from DW-MR acquisitions using standard models like NODDI [21] or SMT [22]. In fact, as each streamline represent a coherent set of real anatomical fibers, there cannot be space for every possible reconstructed streamline. To implement the second prior, we first grouped together all streamlines connecting the same pairs of gray-matter regions and rearranged the corresponding columns of **A** accordingly, as shown in Fig. 2. Then we added a new term to the cost function in Eq. 2 to try explaining the data, if possible, using the minimum number of such groups. Mathematically this is is achieved with the *group lasso* regularization [23] and the problem 2 becomes:

**Figure 2:**
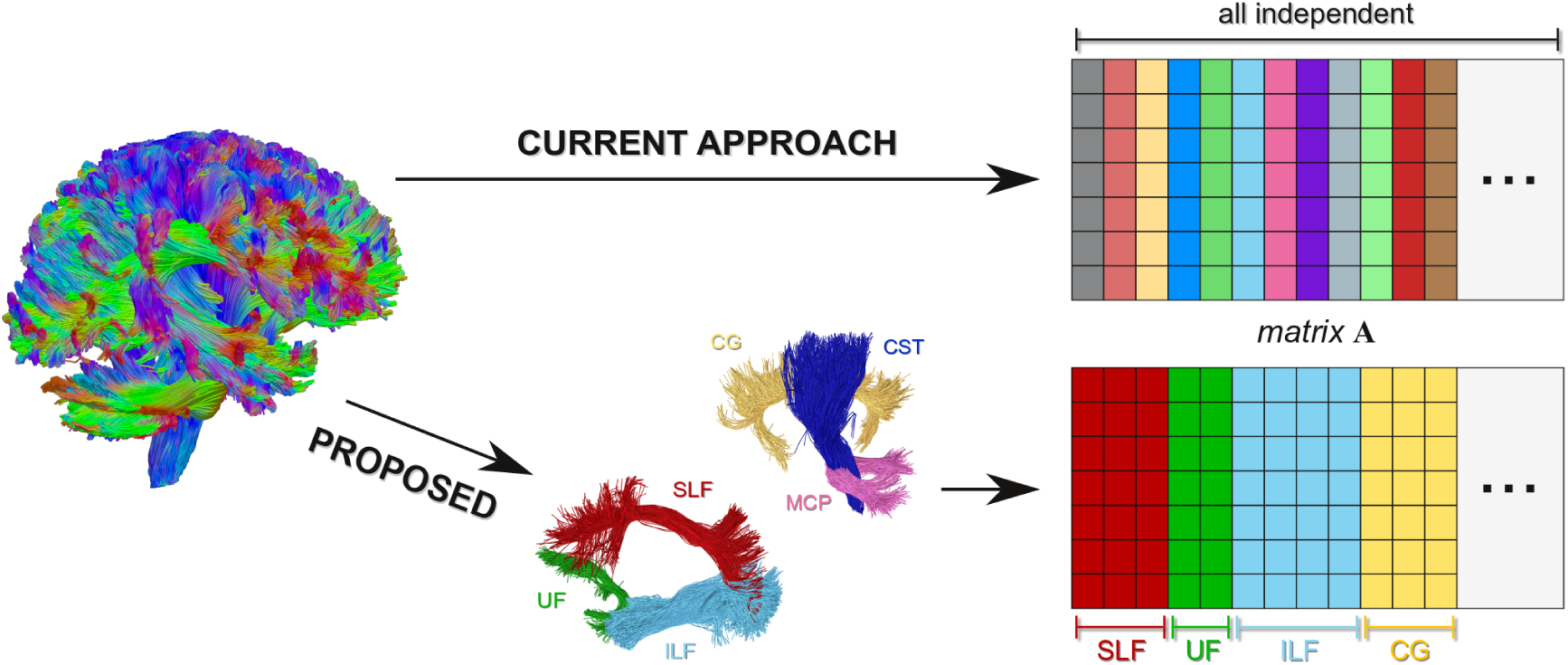
Current tractography algorithms consider all streamlines in a tractogram as independent entities and COMMIT is no exception; every column of the matrix **A** encodes a different streamline and all columns are treated as independent during the estimation of their contributions (top right). The proposed method (COMMIT2) groups streamlines belonging to the same anatomical bundle together and considers the corresponding columns of **A** as a single entity in the estimation; every streamline is still modeled by a distinct column but streamlines of the same bundle are arranged together as a sub-block of the matrix and considered as a whole (bottom left).

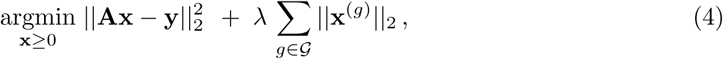

where 𝒢 is a general partition of the streamlines into groups, **x**^(*g*)^ represent the coefficients corresponding to the streamlines in a given group *g ∈ 𝒢* and the parameter *λ >* 0 controls the trade-off between data and the regularization term. This additional term in the cost function penalizes the contributions at the level of groups and, in practice, promotes (but does not constrain) convergence towards a solution that explains the signal with the minimum number of bundles. Note that setting *λ* = 0 corresponds to the classical COMMIT. As this formulation represents an extension to the COMMIT framework, we will refer to it as COMMIT2 in the remaining of the manuscript.

Without any strong a priori knowledge on the bundles, a classical way to operatively solve this problem is to use the so called *adaptive group lasso* [24] which penalizes all groups in the same manner independently of their cardinality. The problem can then be rewritten as:

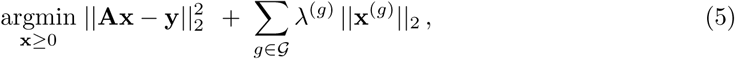

with

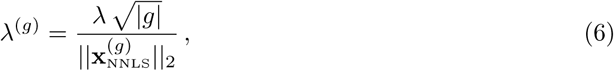

where *|g|* is the cardinality of the group *g* and 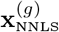 are the weights of the streamlines obtained by solving the NNLS problem in Eq. 2, i.e. without any regularization term.

### 2.3 Data and experiments

We quantitatively evaluated our novel approach using the synthetic phantom developed for the Reconstruction Challenge organized in 2013 at the IEEE International Symposium on Biomedical Imaging [25]. This simulated dataset is shown in Fig. 3a and consists of 27 ground-truth fiber bundles that were specifically designed to mimic real fiber configurations typically encountered in the brain (Fig. 3b). These include complex arrangements of bending, crossing and branching fibers, at various angles and with different curvatures; in addition, three spherical regions corresponding to fast diffusive compartments such as in brain ventricles were added. The intra-axonal signal fraction of this phantom was computed from the geometry of the ground-truth streamlines. The corresponding DW-MR signal was generated using the Composite Hindered And Restricted Model of Diffusion [26] along 64 directions with *b* = 3000s/mm^2^ and adding Rician noise with a signal-to-noise ratio (SNR) of 30. Fig. 3c shows the white-matter mask used for tractography as well as the 53 regions of interest (ROIs) that define the nodes of the corresponding connectome. Figg. 3d-e illustrate two examples of true-positive and false-positive bundles that may potentially be reconstructed with any tractography algorithm. The ground-truth connectivity of this dataset is shown in Fig. 3f and is represented as a graph: the 53 ROIs are displayed as blue circles that are connected by green or red arcs depending whether there is a true-positive and false-positive bundle of streamlines between them, respectively.

**Figure 3:**
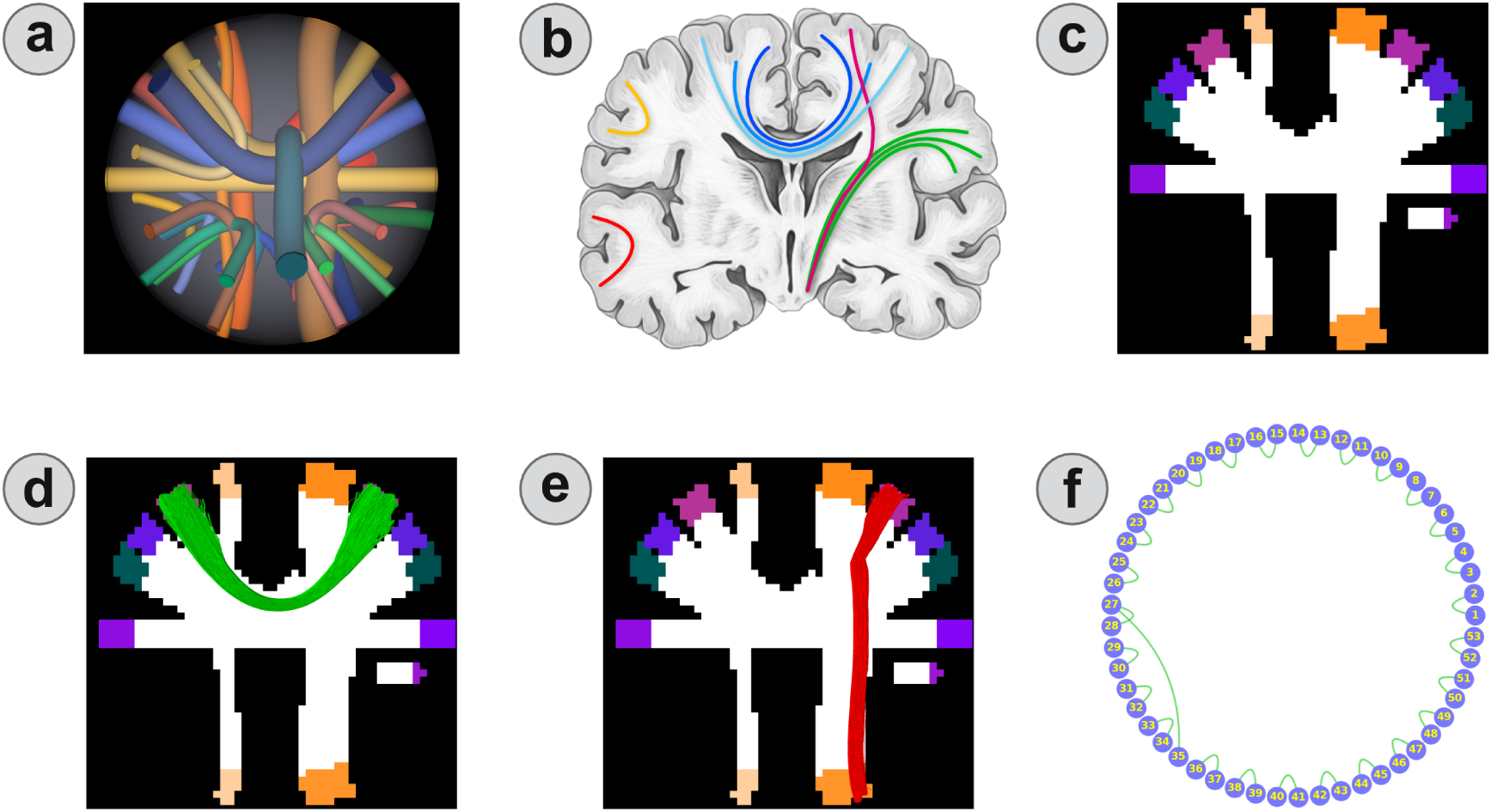
Synthetic dataset used to quantitatively evaluate our method (**a**), inspired from real anatomical bundles of the human brain (**b**). White-matter and gray-matter masks used for tractography (**c**). Examples of true-positive (**d**) and false-positive (**e**) bundles that can potentially be reconstructed with tractography. Ground-truth connectivity represented as a graph (**f**): blue circles correspond to the 53 gray-matter ROIs of (**c**), whereas green and red arcs represent true-positive and false-positive bundles, respectively; please note that no false positives are present in the ground truth.

#### Connectome estimation

Connectomes were constructed from the reconstructions obtained with both *deterministic and probabilistic tractography* using the 53 gray-matter ROIs as network nodes. We employed the MRtrix software [27] as it is a very popular processing suite to analyze DW-MR data. First, we computed the fiber orientation distributions (FOD) in each voxel using Constrained Spherical Deconvolution [28] with *ℓ*_*max*_ = 8. Then, we reconstructed 1 million streamlines with both deterministic (SD_STREAM) and probabilistic (iFOD2) algorithms, using default parameters, and performing the tracking using the WM mask as seed region. Finally, we assigned each endpoint of a streamline to a node if that point fell within 2 mm from one of the 53 gray-matter ROIs (default setting); a streamline was considered as *connecting* two nodes if both endpoints were assigned, otherwise it was discarded and excluded from the analysis.

#### Evaluation criteria

We assessed the sensitivity and specificity of a connectome using the Tractometer metrics defined in [29]. True positives are described in terms of the *Valid Connections* (VC) ratio, which is the proportion of streamlines in the tractogram that connect a correct pair of ROIs, as well as the corresponding number of *Valid Bundles* (VB). Similar metrics can be computed for the false positives, i.e. *Invalid Connections* (IC) and *Invalid Bundles* (IB). To summarize sensitivity and specificity in a single score, we computed the *Youden’s index J* = sensitivity + specificity − 1. Sensitivity is defined as the ratio between VB and the number of real positive bundles (27 in this dataset), and specificity as 1-IB/N, where N is the number of real negatives (462 in this dataset). N represents the number of ROI pairs that may potentially be connected (incorrectly) by tractography and was computed by reconstructing 10 million streamlines with the probabilistic algorithm, for it is more permissive.

## 3 Results

Fig. 4 analyzes the quality of the reconstructions that can be obtained by processing the raw tractogram using the proposed method. These plots correspond to probabilistic tracking, but similar results are obtained with the deterministic algorithm. In the first row are reported the number of valid and invalid bundles (VB and IB respectively) as a function of the regularization strength *λ*. Values at *λ* = 0 correspond to the raw/unprocessed tractogram: VB=27 (corresponding to a sensitivity of 100%) and IB=393 (specificity 14.9%). We can notice that, as *λ* increases, the number of IB decreases rather quickly but, correspondingly, the VB exhibit a much slower decrease trend. The decreasing rate of the IB slows down when they 65reach a value comparable with the VB. However, as expected, by increasing the regularization even further the number of VB also begins to decrease because, as it is known in optimization theory, when *λ* is too large the second term of Eq. 4 dominates and all groups are progressively discarded. To help choosing the optimal value for *λ* we made use of the Youden’s index (J), which is shown in the second row along with the percentage of valid connections (VC). The maximum value of J is about 0.96, which corresponds to VB=27 (sensitivity 100%) and IB=20 (specificity 95.7%). After it reaches its maximum, J starts decreasing as more and more VB are suppressed. Nonetheless, it is interesting to note that the percentage of VC exhibits an increasing trend also after this value, indicating that the rate of decrease of the IB if faster than the VB.

**Figure 4:**
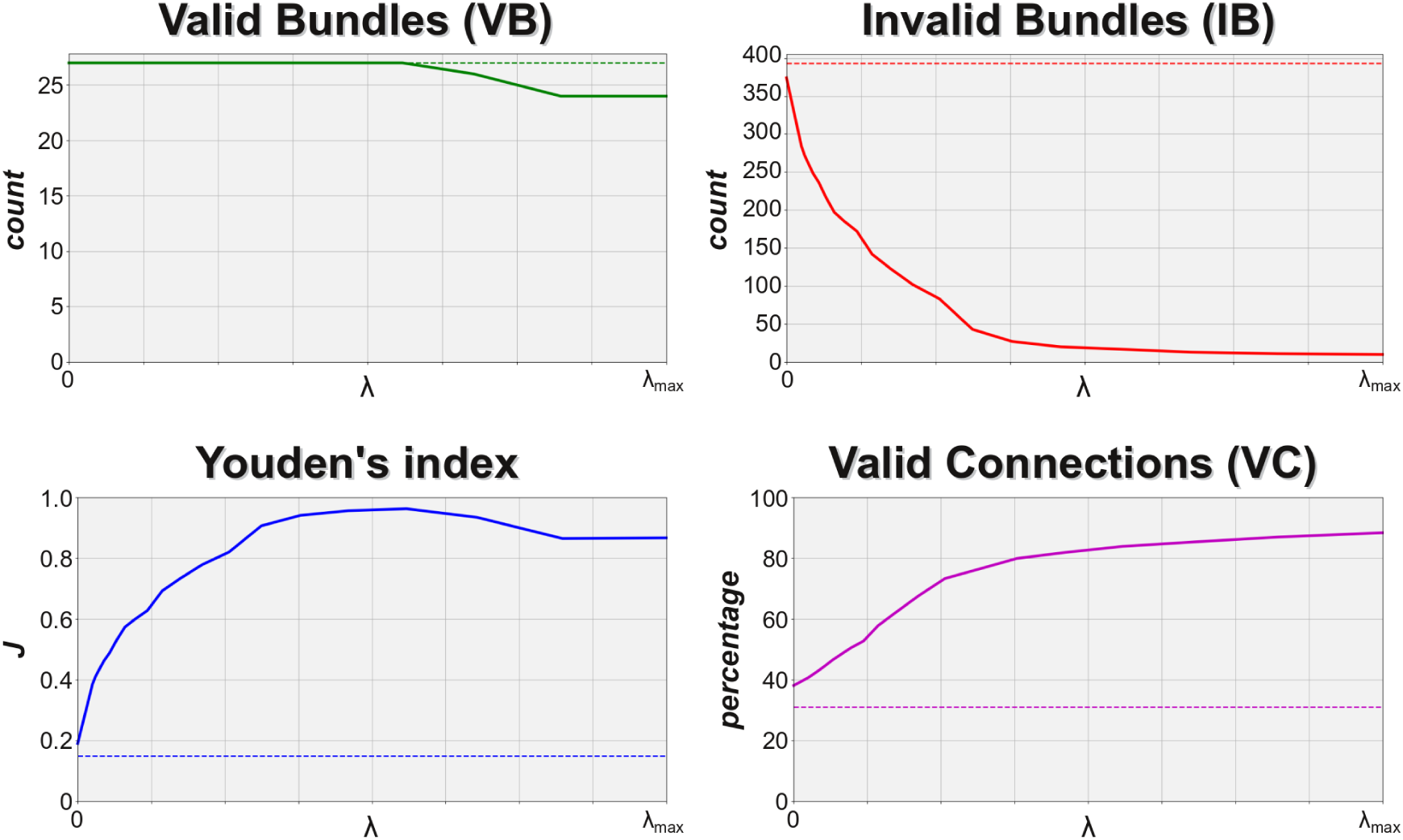
Impact of adding the bundle-wise anatomical priors to COMMIT on the quality of the reconstructions. In the first row are reported the number of valid (VB) and invalid (IB) bundles as function of the regularization strength (*λ*). In the second row are reported the Youden’s index and the percentage of valid connections (VC). Results correspond to probabilistic tractography.

Fig. 5 reports the number of valid bundles (VB) and invalid bundles (IB) bundles in the raw tractogram, as well as in the ones filtered with COMMIT and COMMIT2. Results are shown for both probabilistic and deterministic tractography and correspond to the regularization parameter *λ* that maximizes the Youden’s index J. Both tracking algorithms were able to reconstruct all 27 true bundles, i.e. high sensitivity, but at the price of recovering a large amount of false positives, i.e. very low specificity (IB=393 in case of probabilistic tracking and IB=204 for deterministic). These results agree with previous literature [9–11]. We can clearly see that when COMMIT or COMMIT2 are applied the sensitivity is not affected, as both tractograms still contain all 27 true bundles, while the inclusion of anatomical priors has a dramatic impact on the specificity. Using COMMIT (i.e. no anatomical priors, second column) the number of IB diminished only marginally (393 *→* 374 and 204 *→* 190, respectively). On the other hand, the last column clearly shows that when using COMMIT2 the number of IB is dramatically reduced (393 *→* 20 and 204 *→* 17, respectively).

**Figure 5:**
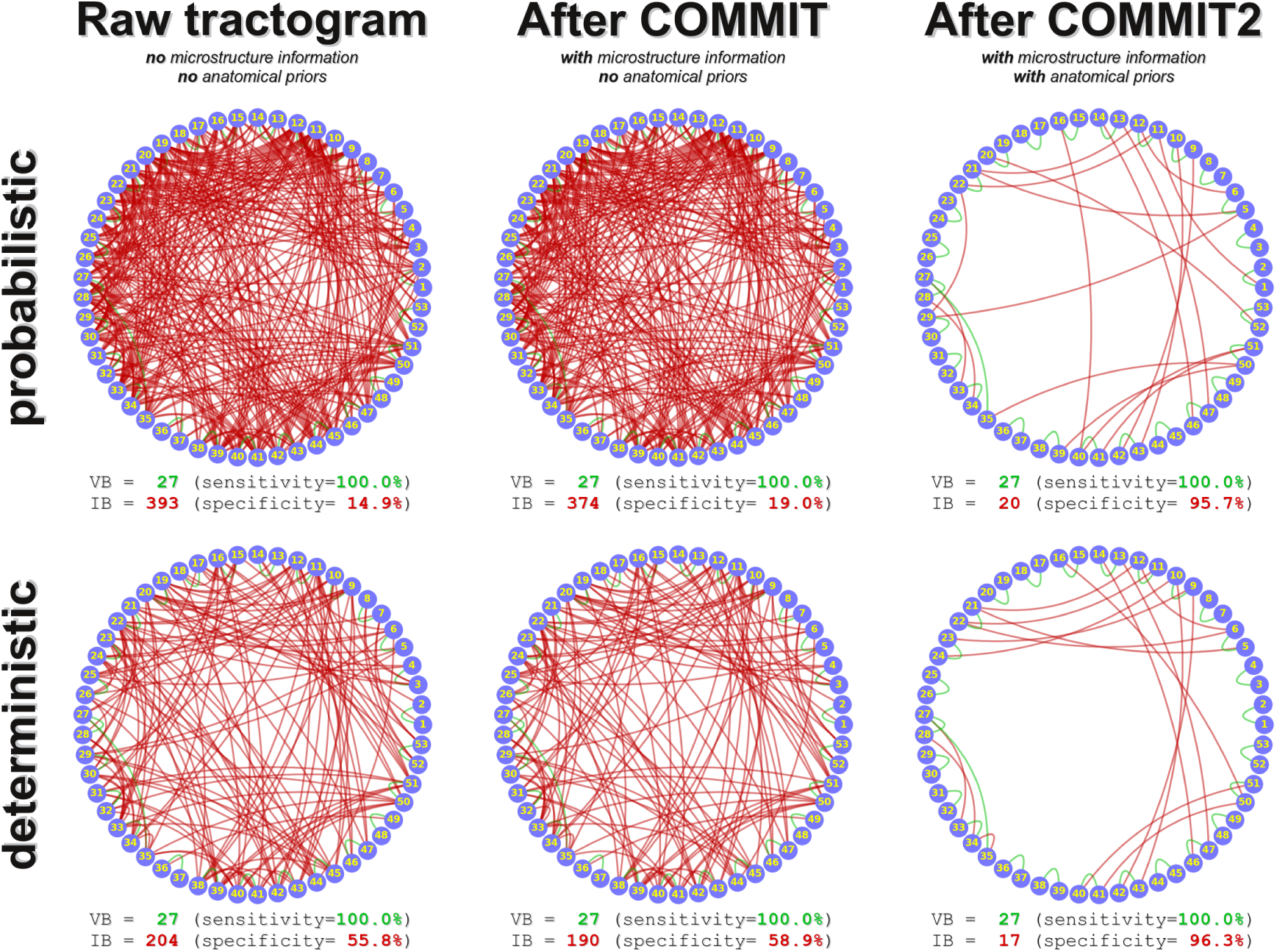
Sensitivity and specificity of tractography before (first column) and after applying COMMIT and COMMIT2 (second and third column, respectively). In all subplots, we identify the true positives in “green” and the false positives in “red”. When using probabilistic tractography (first row) the raw tractogram contained VB=27 and IB=393; note that the true positives are barely visible. After COMMIT, the IB were slightly reduced (from 393 to 374), whereas with COMMIT2 their number was dramatically decreased from 393 to 20. In all cases all 27 true bundles were preserved. Deterministic tractography led to similar results (second row): the IB were reduced from 204 in the raw tractogram to 190 with COMMIT and 17 with COMMIT2. Results hold also for other combinations of tracking algorithm and parameters.

## 4 Discussion

It is worth to highlight that, since the proposed framework allows us to directly inject priors on the bundle, we could also set these values of *λ*^(*g*)^ in terms of known anatomical bundles. In practice, if we are sure that a connection surely exists (from a population study or any atlas), we can promote the corresponding group by setting *λ*^(*g*)^ = 0. On the contrary, if we know that a chosen tractography algorithm is keen to find a particular implausible connection, we can set the corresponding *λ*^(*g*)^ to a very high number which translates in penalizing this group more than the others in the connectome. Another way to take advantage of this property of COMMIT2 is to inject priors coming from other imaging modalities that provide information on bundles. For example, we could set the regularization parameters in terms of functional connectivity results obtained by analyzing functional MRI or magnetoencephalography data that provide us which grey matter ROIs are functionally connected (directly or indirectly).

To explain how COMMIT works with the toy example in Fig. 1, we used the simple 2-compartment “ball and stick model” [20] to build the operator **A**. This choice was made to not burdening the notations and make the exposition as easy as possible. Nevertheless, it is worth noticing that the flexibility of COMMIT (and then COMMIT2) allow us to use any microstructure model to describe the DW-MR signal (as presented in the original papers [16, 17]). In accordance with the acquisition parameters, we can thus take more complicate models whose performances have been already investigated in literature. For example, in this work we showed that COMMIT2 framework can be used to fit not only the entire DW-MR signal acquired, but also to any microstructure map (derived by any imaging modality) provided that the microstructure feature to be decoupled on each streamline can be considered as invariant along the path. For example, considering the resolution we have with MRI images, we in this work we supposed that the intra-axonal sectional area could be considered to be constant for each streamline, since it represents the intra-axonal sectional area of a group of axons sharing the same path. This property of COMMIT opens to the possibility of using multimodal images also to assign contributions to the streamlines and not only to penalize the bundles as mentioned before. A valid option could be to use myelin maps derived from MR relaxometry, quantitative magnetization transfer or quantitative susceptibility. By using COMMIT2 to fit simultaneously an intra-axonal and a myelin signal fraction maps, one could estimate at the same time the intra-axonal and the myelin volumes associated to each individual streamline and thus compute pathway specific g-ratio.

### 4.1 Limitations and future perspectives

Although we obtained outstanding results, we acknowledge that the proposed framework is not without limitations and there is room for future improvements. First, the bundle regularization guarantees that if a bundle is necessary to explain the signal, then all its streamlines will be kept. This implies that none of the eventual redundant streamlines present in a group will be eliminated; rather the weights will be equally distributed among them resulting in very small contributions. Moreover, even if a streamline follows a very different path from the other in the same group, because of this choice of regularization, it will kept since it still connects the same two ROIs. Although a proper way to filter inside the groups is yet under investigation and will be object of future works, we can speculate that one possible way to do that is considering a finer parcellation for the gray matter resulting in smaller groups to be evaluated by the framework. Another way could be using clustering techniques to group streamlines together (e.g. [30, 31]). All these possibilities will be tested and compared in future analysis.

Another current limitation is the choice of the parameter *λ* that scales the groups penalization. From the literature we know that if the columns of the operator **A** are linearly independent there exists an upper (*λ*_*max*_) an lower (*λ*_*min*_) bounds for *λ* (see [24] as reference). However, this is not the case for a general tractogram, because the same (or geometrically equivalent) pathway could be shared by more than one streamline inducing redundancy inside **A**. In this work we set *λ*_*min*_ = 0, which provided the results of the standard COMMIT framework, and we found the *λ*_*max*_ empirically evaluating the loss of fibers and stopping when too many bundles were discarded resulting also in a worse fit. Future investigations to possibly set a priori a *λ*_*max*_, and consequently find the optimum *λ* are needed especially when using COMMIT on in vivo data.

## 5 Conclusion

Recent studies have shown that current tractography algorithms are quite good at reconstructing the major WM bundles, i.e. high sensitivity, but at the price of also recovering a large amount of false positive ones, i.e. low specificity. In particular, those false positive streamlines can drastically bias any subsequent analysis, as non-existent structures are mixed with real ones. In this work, we showed that adding basic prior information on the organization of the neuronal connections can help microstructure informed tractography in dramatically improving the quality of reconstructions. Moreover, our novel formulation (COMMIT2) opens up the possibility of injecting any prior knowledge of the bundles in the framework. This leads to the opportunity of including multimodal acquisitions in a unique framework resulting in the reconstruction of a more veridical connectome. In conclusion, our results represent an important step forward to boost the accuracy of tractography and may have profound implications for the use of tractography to study structural brain connectivity.

## Acknowledgements

This work was supported by the Rita Levi Montalcini Programme for young researchers of the Italian Ministry of Education, University and Research (MIUR).

